# Systemic pesticides in a solitary bee pollen food store affect larval development and increase pupal mortality

**DOI:** 10.1101/2022.05.12.491686

**Authors:** Ngoc T. Phan, Neelendra K. Joshi, Edwin G. Rajotte, Fang Zhu, Kari A. Peter, Margarita M. López-Uribe, David J. Biddinger

## Abstract

Solitary bees are often exposed to various pesticides applied for pest control on farmland while providing pollination services to food crops^1^. Increasing evidence suggests that sublethal toxicity of agricultural pesticides affects solitary bees differently than the social bees used to determine regulatory thresholds like honey bees and bumblebees^2–4^. Studies on solitary bees are challenging because of the difficulties in obtaining large numbers of eggs or young larvae for bioassays. Here we show the toxic and sublethal developmental effects of four widely used plant systemic pesticides on the Japanese orchard bee (*Osmia cornifrons*). Pollen food stores of this solitary bee were treated with different concentrations of three insecticides (acetamiprid, flonicamid, and sulfoxaflor) and a fungicide (dodine). Eggs were transplanted to the treated pollen and larvae were allowed to feed on the pollen stores after egg hatch. The effects of chronic ingestion of contaminated pollen were measured until adult eclosion. This year-long study revealed that chronic exposure to all tested pesticides delayed larval development and lowered larval and adult body weights. Additionally, exposure to the systemic fungicide resulted in abnormal larval defecation and increased mortality at the pupal stage, indicating potential risk to bees from fungicide exposure. These findings demonstrate potential threats to solitary bees from systemic insecticides and fungicides and will help making policy decisions for mitigating these effects.

## Main

Pollination is a vital process in agricultural crop production. Bees contribute to the abundance and variety of global food offerings and the great economic value of agricultural commodities^5^. Without bees, some food crop yields would be reduced by up to 90%^6^. The solitary mason bees (*Osmia* spp.) have been widely used for crop pollination in Asia, Europe, and North America^7–9^. They are short-ranged pollinators that tend to stay in the pollination-targeted crop and can be reared in large numbers to commercially pollinate fruit crops making them suitable for pesticide toxicity evaluations representing non-Apidae bees^10–12^. *Osmia* spp. are univoltine, solitary bees that fly for only a few weeks in the early spring. Female *Osmia* nest in abandoned beetle holes in dead trees or hollow stems as well as cardboard or wooden tubes when commercially managed for crop pollination. Pollen is carried on the female abdominal scopa and packed into round pellets inside individual cells, which are arranged in a series partitioned by mud walls. The pollen pellet is the food provision (store) for a single larva^8^. Unlike honey bee larvae, which are taken care of by nurse bees from egg hatch until pupation and mainly fed by glandular secretions, *Osmia* larvae are not tended by adults and feed solely on the single pollen provision during the summer. Pupation occurs within the cell in the fall and eclosion to overwintering adults quickly follows, and adults emerge the following spring^13^.

One of the major threats that bees face is exposure to agricultural pesticides during their pollination activities on crops^13^ that could be lethal alone or cause increased bee mortality by synergistically interacting with other stressors such as diseases, parasitic mites, and additional pesticides^14^. These other pesticides include insecticides, fungicides, and other agrochemicals that are necessary to maximize production and protect crops against pests. Pesticides are generally applied as foliar sprays, soil applications, or seed treatments^15^. These pesticides can be restricted to plant surfaces relying on contact toxicity to the pest, or they can be absorbed into the plant and by either translaminar movement (locally systemic) or transported throughout the plant via xylem and/or phloem tissues to reach all plant tissues (truly systemic). Even systemic pesticides can also have varying levels of contact activity soon after application, but this activity usually disappears within a few days as the surface residues are broken down by sunlight or weathered in other ways. Systemic pesticides confer some level of protection to useful arthropods that do not feed on the plants such as biological control agents. However, they can be translocated to pollen and nectar via either xylem or phloem, although usually at much lower levels than the initial application rates^16^. During bloom, bees collect pollen and nectar as the main source of protein, lipids, and energy. Pesticide contaminated food collected and used as nest provisions may affect bee offspring. Since *Osmia* larvae live exclusively on pollen provisions, pesticide exposure may seriously affect development and reduce the next season’s bee population. Furthermore, *Osmia* bees are univoltine, and their populations do not recover from pesticide poisoning within the same season as multivoltine social bees (e.g., honey bees and bumble bees)^17^.

In this study, we examined the effects of three commonly used systemic insecticides and one fungicide from egg through adult on the development and survivorship of the Japanese orchard bee, *Osmia cornifrons* (Hymenoptera: Megachilidae). This is a solitary mason bee that has been managed for fruit pollination in the eastern U.S. and Japan^18,19^ and is amenable to produce the large numbers of bees necessary for conducting pesticide bioassays necessary for regulatory risk assessment. However, there is no information on how different pesticides directly affect the developmental stages of this species. Therefore in this context, we conducted a year-long study using formulated pesticide products and examined the Low (1/10X), Medium (1X), and High (10X) concentration exposure based on their field-realistic concentrations (1X) specified in a previous study^16^. In that study, field-realistic pesticide concentrations in nectar and pollen were measured in apple orchards that were sprayed using grower application equipment and practices five days before apple bloom (pink bud developmental stage). Using a modified natural pollen provision feeding method^20^, we assessed the effects of pesticide exposures on larval development times, last larval instar body weight, and survivorship through adult eclosion. This study will help in understanding the risks that systemic pesticides including fungicide can pose to the development of solitary bees from pre-bloom and bloom pesticide applications in various crops.

Overall, the survival rate of *O. cornifrons* larvae was 100% for all pesticides at 1/10X (low) and 1X (medium) concentrations except for the fungicide dodine where survival was significantly reduced to 89.83%. At the highest (10X) rate, survival was also significantly reduced for larvae exposed to acetamiprid, sulfoxaflor, and dodine pesticides (F_12, 65_ = 7.02, *p* < 0.001) (Table 1). However, sublethal effects of pesticides on larvae caused extended pre-pupal development time, and low larval weight. In addition, the high concentration (10X rate) of fungicide dodine via pollen food stores caused abnormal defecation by larvae (F_12, 65_ = 115.24, *p* < 0.001). Under this treatment, the larval meconium of 66.83% larvae was in long strands rather than normal fragmented frass (Table 1, Fig. 1). The average time to the cocoon spinning stage was 20.44 days in the control group. All larvae exposed to pesticides took significantly longer to reach the cocoon spinning (pre-pupae) stage than the control, even those in the lowest 1/10X rate e (F_12, 611_ = 28.80, *p* < 0.001) (Table 1). Within the 1/10X rate group of treatments, larval development was delayed by 2.56– 3.37 days compared to the control, but none of the pesticides were significantly different from each other. Within the 1X rate group, larval development was delayed by 2.64–3.98 days compared to the control with only the sulfoxaflor 1X treatment delaying development significantly longer than those of the 1/10X rate group. Within the 10X rate group, larval development was delayed by 3.02–4.35 days with both the sulfoxaflor and dodine treatments resulting in significantly longer delays than the lowest (1/10X) rate group. Of the five pesticides, acetamiprid and flonicamid insecticides caused the least delay in larval development. In the fungicide treatments, the 10X rate of dodine caused the longest developmental delay of 4.35 days, which was significantly longer than both the other rates.

**Table 1:**
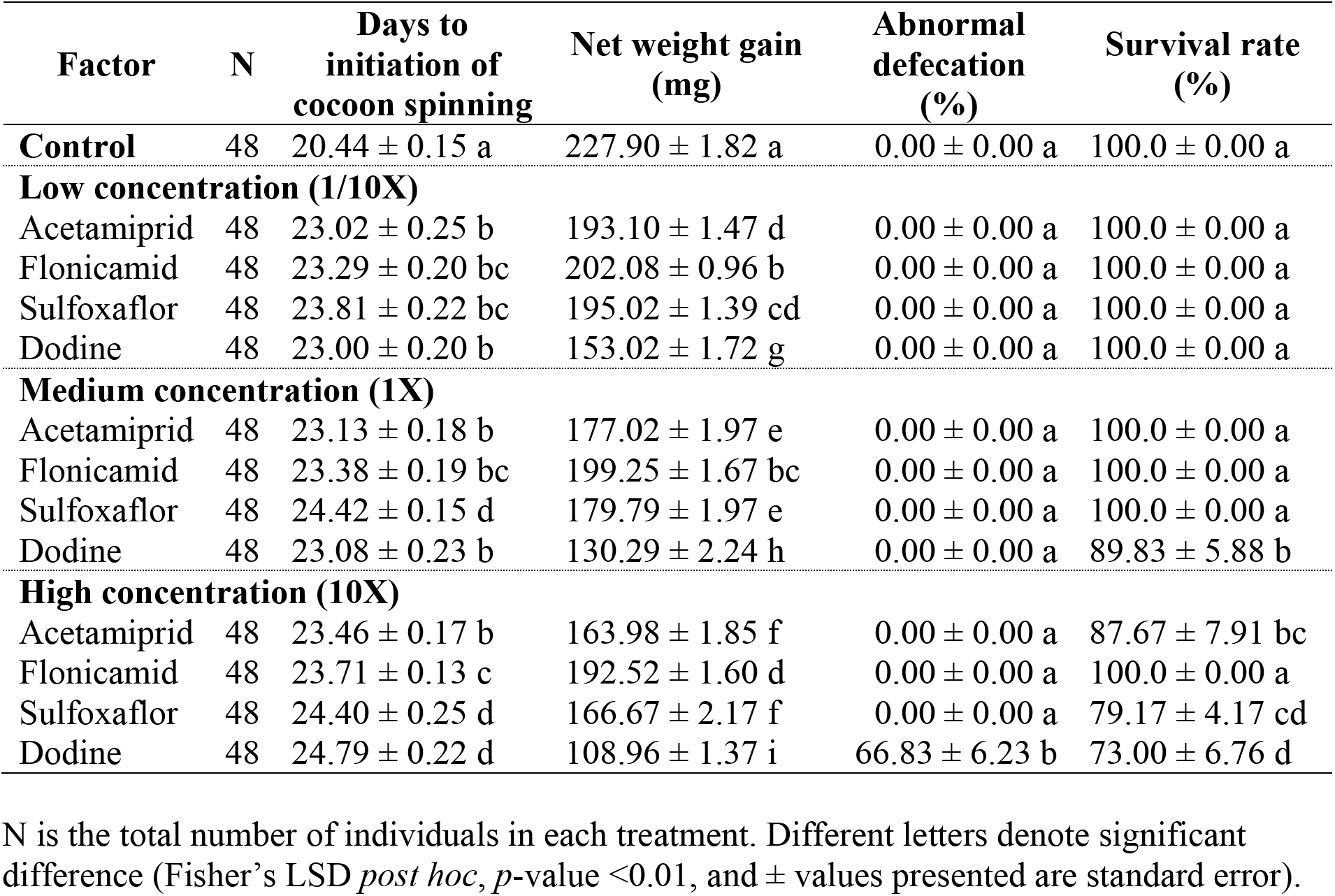
Effect of different treatments and exposure levels of selected pesticides on development time, net weight gain of 5^th^ instar, and survivorship to adult of the Japanese orchard bee larvae.

| Factor | N | Days to initiation of cocoon spinning | Net weight gain (mg) | Abnormal defecation (%) | Survival rate (%) |
| --- | --- | --- | --- | --- | --- |
| **Control** | 48 | 20.44 ± 0.15 a | 227.90 ± 1.82 a | 0.00 ± 0.00 a | 100.0 ± 0.00 a |
| **Low concentration (1/10X)** |  |  |  |  |  |
| Acetamiprid | 48 | 23.02 ± 0.25 b | 193.10 ± 1.47 d | 0.00 ± 0.00 a | 100.0 ± 0.00 a |
| Flonicamid | 48 | 23.29 ± 0.20 bc | 202.08 ± 0.96 b | 0.00 ± 0.00 a | 100.0 ± 0.00 a |
| Sulfoxaflor | 48 | 23.81 ± 0.22 bc | 195.02 ± 1.39 cd | 0.00 ± 0.00 a | 100.0 ± 0.00 a |
| Dodine | 48 | 23.00 ± 0.20 b | 153.02 ± 1.72 g | 0.00 ± 0.00 a | 100.0 ± 0.00 a |
| **Medium concentration (1X)** |  |  |  |  |  |
| Acetamiprid | 48 | 23.13 ± 0.18 b | 177.02 ± 1.97 e | 0.00 ± 0.00 a | 100.0 ± 0.00 a |
| Flonicamid | 48 | 23.38 ± 0.19 bc | 199.25 ± 1.67 bc | 0.00 ± 0.00 a | 100.0 ± 0.00 a |
| Sulfoxaflor | 48 | 24.42 ± 0.15 d | 179.79 ± 1.97 e | 0.00 ± 0.00 a | 100.0 ± 0.00 a |
| Dodine | 48 | 23.08 ± 0.23 b | 130.29 ± 2.24 h | 0.00 ± 0.00 a | 89.83 ± 5.88 b |
| **High concentration (10X)** |  |  |  |  |  |
| Acetamiprid | 48 | 23.46 ± 0.17 b | 163.98 ± 1.85 f | 0.00 ± 0.00 a | 87.67 ± 7.91 bc |
| Flonicamid | 48 | 23.71 ± 0.13 c | 192.52 ± 1.60 d | 0.00 ± 0.00 a | 100.0 ± 0.00 a |
| Sulfoxaflor | 48 | 24.40 ± 0.25 d | 166.67 ± 2.17 f | 0.00 ± 0.00 a | 79.17 ± 4.17 cd |
| Dodine | 48 | 24.79 ± 0.22 d | 108.96 ± 1.37 i | 66.83 ± 6.23 b | 73.00 ± 6.76 d |

N is the total number of individuals in each treatment. Different letters denote significant difference (Fisher's LSD *post hoc*, *p*-value <0.01, and ± values presented are standard error).

**Figure 1.**
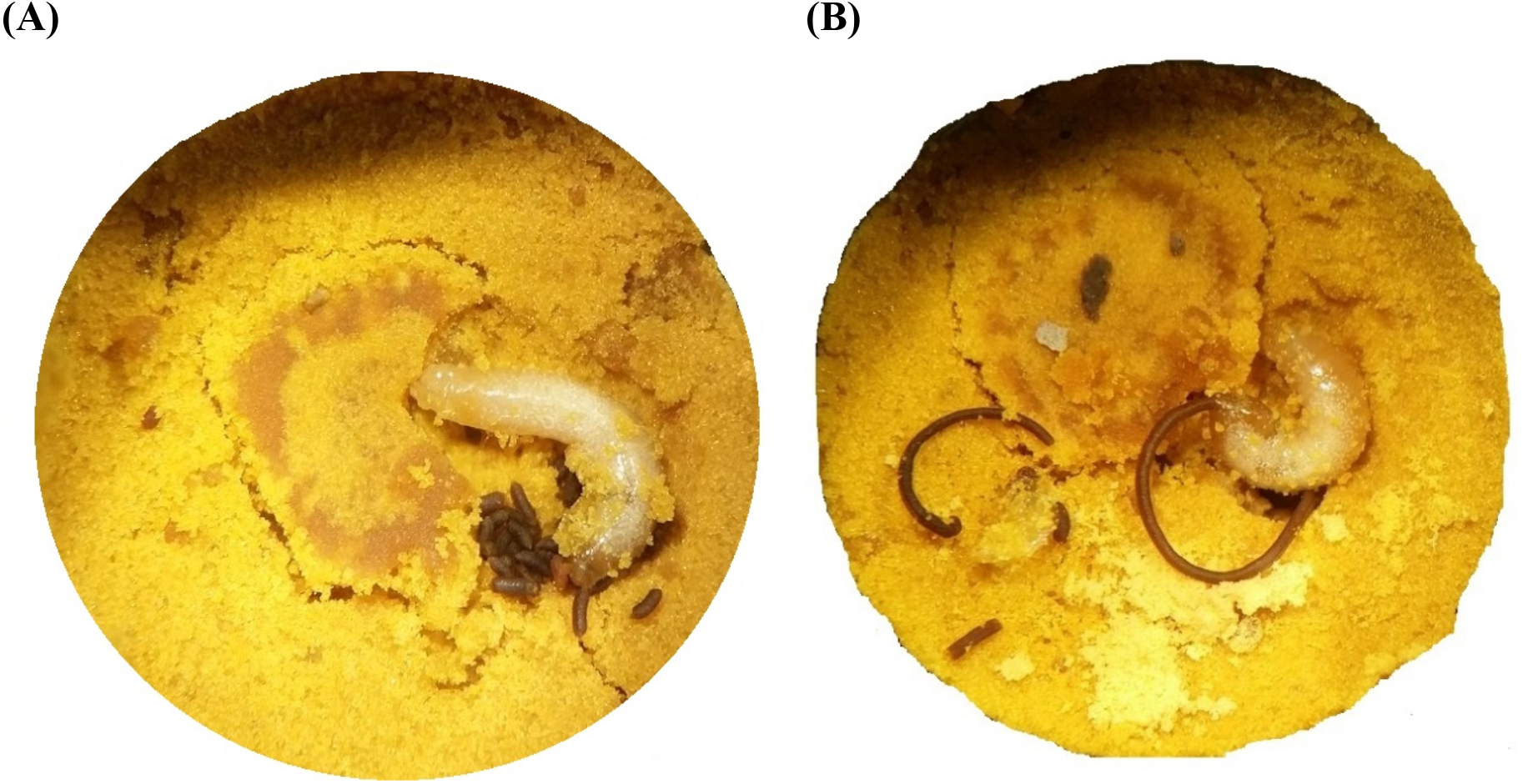
Japanese orchard bee larvae (5^th^ instar) with their frass. (A) Normal defecation; (B) Abnormal defecation.

All pesticide treatments caused lower net weight gains in larvae than the control and were inversely related to exposure level (F_12, 611_ = 334.89, *p* < 0.001) (Table 1). Flonicamid insecticide had the lowest reduction in weight gain of all treatments (11.33–15.52%) with the 10X rate being significantly lower than the other two rates. Exposure to acetamiprid and sulfoxaflor insecticides resulted in statistically greater levels of weight loss in larvae than flonicamid, but were statistically equivalent to each other at all corresponding rates. Dodine caused the greatest weight loss in larvae that ranged from 32.86% at the lowest rate, 42.82% at the 1X rate, and 52.19% at the 10X rate that were all statistically different from each other (Table 1). All treatments had significantly lower rates of weight gain than the control and were inversely related to exposure level (F_12, 611_ = 836.06, *p* < 0.001) (Fig. 2). Flonicamid insecticide had the least reduction in daily rate gain compared to the control (10.45–18.88%) with only the highest 10X rate being worse than the other two rates. Reductions in weight gain in other insecticide treatments were statistically similar (Fig. 2). Exposure to fungicide dodine caused significantly higher losses in the average daily weight gain (28.12–43.41%) than other pesticides and the control (Fig. 2).

**Figure 2.**
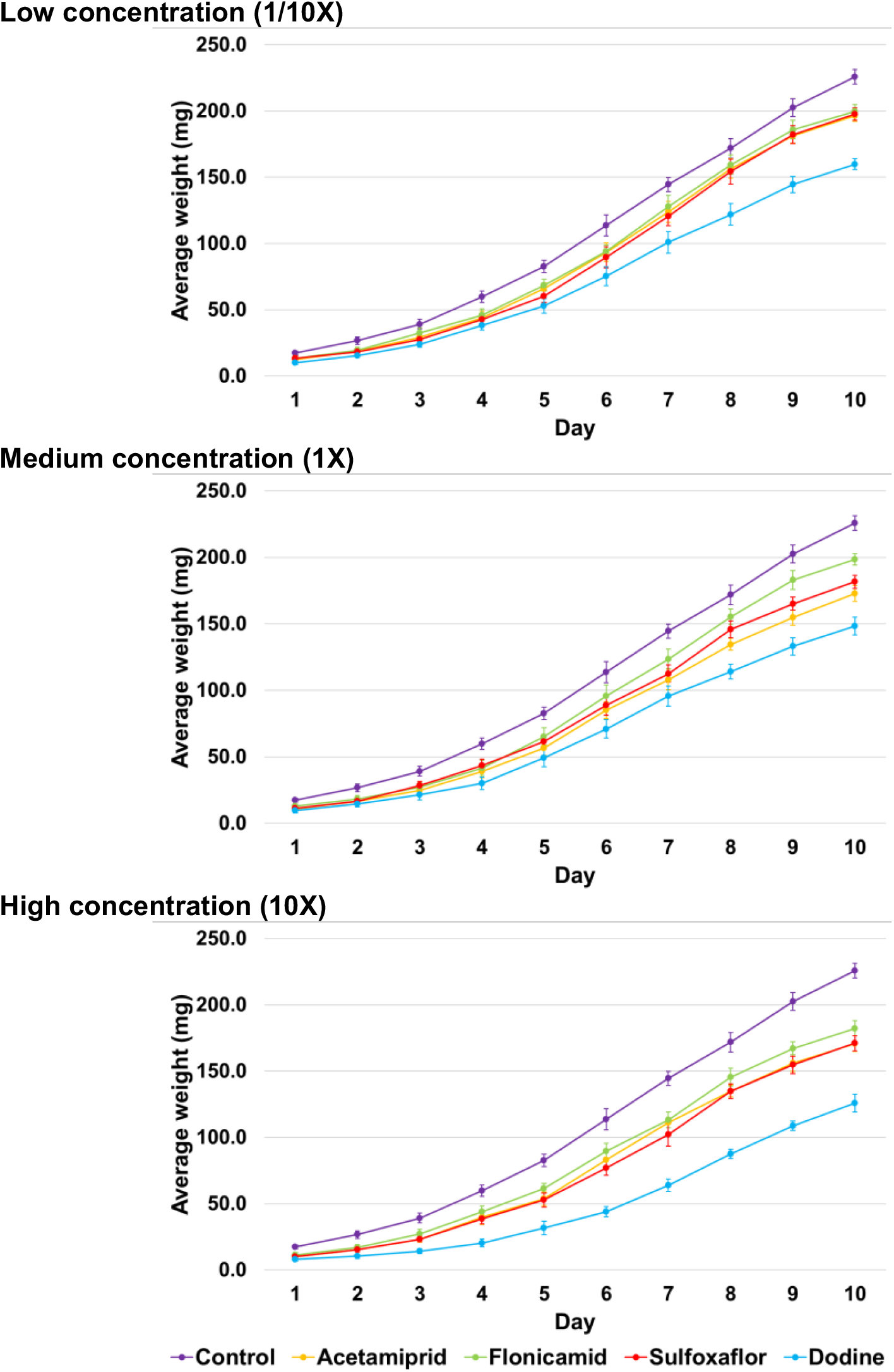
Japanese orchard bee larval weight and weight gain rate for the first 10 days of the 5^th^ instar in different treatments. Dots show the mean and error bars show standard deviation. Lines show the weight gain rate. Different colors represent different pesticide treatments. N = 48 per treatment in each experiment.

Female body weight differs between groups (F_12, 107_ = 209.20, *p* < 0.001) (Fig. 3). All treatments had a significant negative effect on adult body weight and were inversely related to exposure level except for the 1/10X and 1X rates of acetamiprid and all rates of flonicamid, which were not significantly different (Fig. 3). Flonicamid had the lowest reduction in body weight of all treatments (7.63–9.09%). Females exposed to 1/10X and 1X rates of acetamiprid had at 10.61– 19.63% reductions in body weight, which were equivalent to flonicamid. However, at the 10X rate, acetamiprid exposure caused higher body weight loss in females than flonicamid. Sulfoxaflor, at 11.05%–19.72% reductions in body weight gave significantly greater levels of body weight loss than flonicamid at all corresponding rates. Dodine fungicide caused significantly higher losses in average body weight (36.01–50.92%) in female bees than all other pesticides (Fig. 3). Male body weight was significantly lower in all treatments than the control (F_12, 452_ = 758.77, *p* < 0.001) (Fig. 3). Within the 1/10X rate treatments, acetamiprid and flonicamid had the lowest reduction in body weight, but the same rate of sulfoxaflor resulted in statistically greater level of body weight loss than the control (Table 1). Within the lowest rate, significantly descending order of body weight loss was flonicamid (11.45%), sulfoxaflor (17.81%), acetamiprid (22.01%), and dodine (37.02%). Within the highest rate, reductions in body weight were similar between acetamiprid (23.45%) and sulfoxaflor (22.21%) (Fig. 3). All rates of dodine caused significantly higher body weight losses (26.19–49.79%) in male bees than all other treatments (Fig. 3).

**Figure 3.**
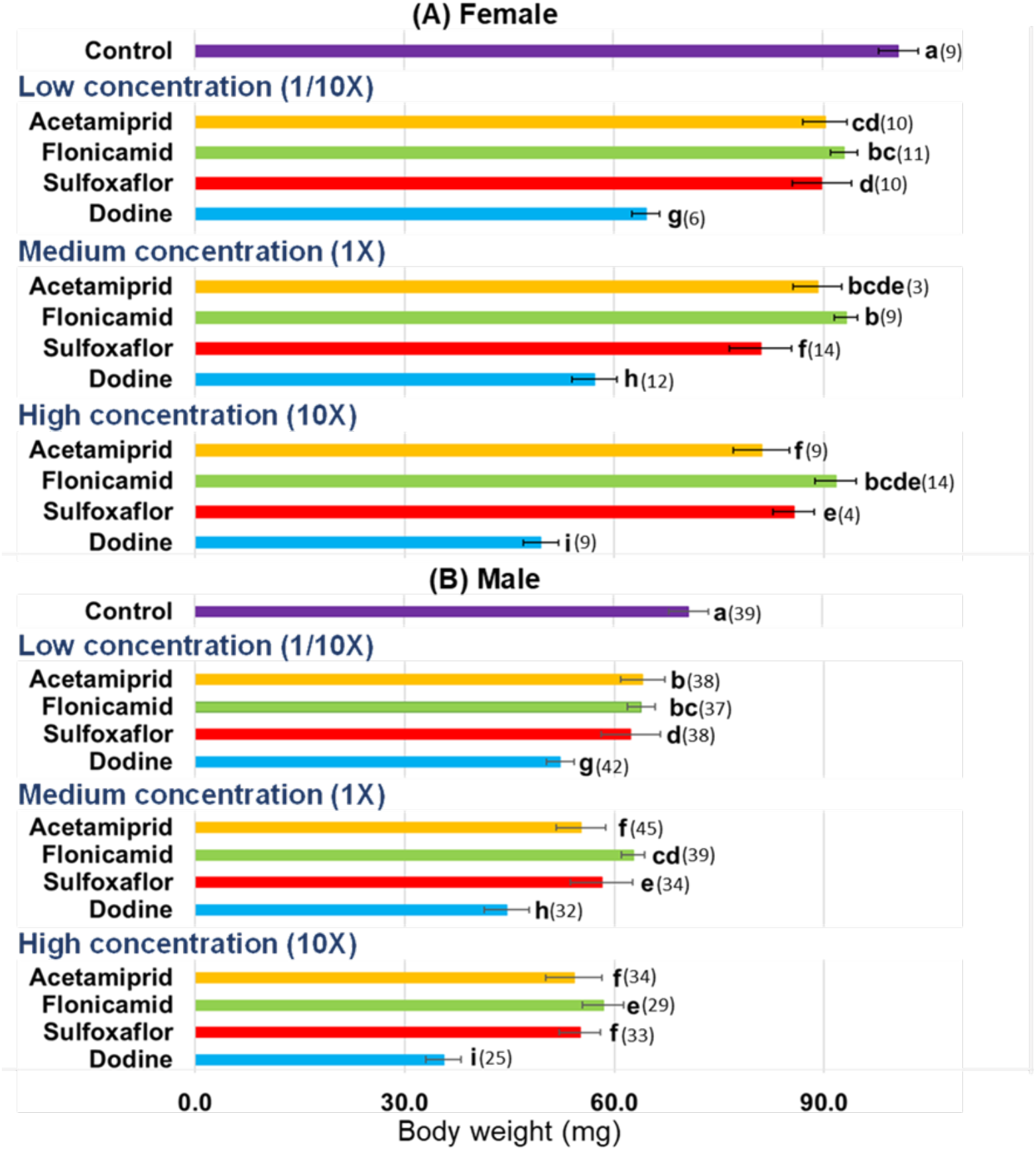
Adult body weight (in mg) of the Japanese orchard bee under exposure in different concentrations of insecticides (acetamiprid, flonicamid, sulfoxaflor) and fungicide (dodine) treatments. Bars show the mean and error bars show standard deviation. Different letters denote significant difference (Fisher’s LSD *post hoc, p*-value < 0.001). Numbers indicate *n* (number of individuals) in each treatment.

This study reveals that the exposure to plant systemic insecticides and a fungicide at field-realistic rates in *O. cornifrons* pollen food stores did not affect larval survival but extended larval developmental time and reduced the daily weight gain of developing last instar larvae. The net weight gain of resulting pre-pupae was also reduced for all treatments, regardless of rate. Lower survival within the pupal stage was found at the 10X rate of all pesticides except for flonicamid and was also found in the 1X rate of the fungicide, dodine. Interestingly, exposure to the fungicide dodine alone caused abnormal larval defecation and significantly lower survival rate. This is the first study documenting impact of a commonly used agricultural fungicide on the development of *O. cornifrons*, which is an efficient pollinator of economically important fruit crops like apple where dodine is used. For social bees, recent studies suggested that consumption of fungicide-contaminated pollen inhibited digestion of worker honey bees *Apis mellifera*, causing malnutrition, which led to reduced population levels. However, larval exposure to fungicide did not affect the development or survivorship to adult^21^. Fungicide impacts were also found in stingless bees *Melipona scutellaris*, causing morphological and physiological changes in their midgut cells, which interrupted nutrition absorption and reduced bee lifespan^22^.

Chronic exposure to the fungicide dodine in the present study also caused decreased survival at the pupal stage, indicating potential threats to bees from systemic fungicide exposure. Fungicides have been sprayed near and during bloom for decades under the assumption that they are not harmful to pollinators^23^. However, synergism between fungicide and other pesticides that are often mixed in the spray tank and applied together, likely occurs in agriculture and can have direct repercussions on bee populations. Recent studies found that fungicide exposure resulted in the downregulation of genes^24–26^, cell death, and the inhibition of P450 detoxification^27^, which led to malnutrition, impaired performance, and increased susceptibility to other pesticides. Several studies suggested that fungicides could negatively affect nesting behaviors and reproductive success of solitary bee species. In bumblebees, the reduction of queen body mass and reproductive capacity due to fungicide exposure led to poor colony performance and decreased bee biomass. In honey bees, exposure to fungicides caused decreased forager survival, impaired hormonal regulation, and behavioral transition in adult bees. Some studies on honey bees even showed that fungicides might affect larval development and reduce adult eclosion^28^. Our study is the first record showing that a fungicide alone could directly cause mortality at the pupal stage of *Osmia*.

Several studies showed that the body size of solitary bees is related to foraging efficiency, homing ability, dispersal, and fecundity^29–33^. In our study, individuals that consumed pollen mixed with pesticides had lower weight gain rates and smaller adult body weight, which suggests further investigation into the chronic impacts of pesticide exposure on the fecundity of surviving adults. We measured adult weight by opening their cocoons and extracting adults to get the maximum weight of adult bees upon the completion of chill requirements necessary to break diapause^34^, but before actual emergence. The natural emergence process of chewing through the cocoon and nest debris in the nest tube could consume a lot of energy that smaller-sized bees with possibly lower lipid reserves may not complete, causing pre-emergence mortality.

Our results show that sublethal effects differ by sex and to examine effects on fecundity or fertility in future studies, targeting and separating eggs of different sexes based on *Osmia* nesting behavior are necessary. Our results also showed that the male bees exposed to different pesticide treatments had significantly lower body weight than the control. Male solitary bees also contribute to pollination and are exposed to pesticides^35^. Therefore, pesticide risk assessment for solitary bees should not just target females because maintaining a sex ratio at 1:1 is essential for reproductive success of these species^36–38^. Since a typical female *O. cornifrons* lays only 30 eggs throughout her life^8^, we do not know how many eggs that a smaller, pesticide-exposed bee can lay and whether a smaller male can ensure reproductive success. Previous studies found that smaller honey bee males had less spermatozoa, were weaker fliers and less successful at mating than normal-sized males^39,40^. Future research addressing these related questions would advance regulatory framework development for pesticide risk assessment for solitary bees. Our findings on the toxicity and sublethal effects of the fungicide dodine alone are important and suggest that further investigations of the impacts of both plant systemic and the more common contact fungicides are warranted for other bee species as well to provide a better understanding of the underlying mechanisms of the toxicity and sublethal effects found in our study.

## Methods

### *Osmia cornifrons* trap nests set-up

Nesting materials, including water-resistant straws (0.307” × 6” Open-Ended Opaque White) and loose cardboard tubes (0.312” × 6” Open-Ended Kraft) (Jonesville Paper Tube Corporation, MI), were kept inside blue-tote shelters (15” × 23” × 16”) (Postal Products Unlimited Inc., WI). Each shelter was placed securely by attaching on top of two wooden crates to avoid being blown away by strong winds. We used chicken wire to cover the front of blue totes to prevent birds from attacking the nests (Fig. S1). These shelters were placed on the mulch strip underneath the apple trees in the Arboretum at Penn State, University Park, PA with their open ends facing Southeast (after Batra 1989)^41^.

*Osmia cornifrons* (sourced from a single supplier in Harleysville, PA) were allowed to emerge in a BugDorm (MegaView Science Co. Ltd., Taiwan) at room temperature (25°C). We kept these adults in the lab for at least three days (up to one week) to ensure mating. An important step before uncaging the bees was to make sure no other bee species (e.g., *O. lignaria* or *O. taurus*) were mixed into our testing population. Adult *O. cornifrons* were then released in the nest shelters in front of the loose tubes. We released a group of ~100 adult bees in the morning every couple of days (depending on the weather) from April 20^th^ to April 30^th^, 2017 to increase the colonization rate in the nest shelters and, most importantly, to extend the egg-laying period for collection. Since mason bees show attraction to aggregation pheromones associated with their cocoons^42^, we kept eclosed cocoons in the shelters in front of the nest blocks to increase nesting success. A single nesting site at the Penn State Arboretum (University Park, PA) was used because it was pesticide-free and had diverse pollen sources. The nests were checked every day and the tubes with pollen provisions were collected. New empty nest tubes were then replaced in the exact location of the collected tubes to prevent the bees from abandoning the nests.

### Preparation of pollen provisions and collection of *Osmia cornifrons* eggs

Artificial pollen provisions of the same average weight were formed in separate clear cells of bio-assay trays with lids (LBS Biotech, United Kingdom) and used for our pesticide bioassays. The average weight of individual *Osmia* collected pollen provisions (290.24 ± 3.55 mg, n=50) was calculated by weighing 50 random unused provisions collected from the nest tubes. Pollen provisions collected from many tubes in the trap nests were homogenized in a mixer (Fig. S2). In pretests, we noted that moisture loss occurred during pollen preparation and that dry provisions were hard to consume by 2^nd^ instar larvae. To maintain proper moisture content, we measured the average moisture content of 50 random provisions with feeding young larvae (2^nd^ instar) collected from the trap nests (25% ± 0.14, n=50). For all bioassays, we subsequently hydrated the homogenized pollen using distilled water to maintain this average of 25% w/w.

Pesticide concentrations were added to ~290mg-artificial pollen provisions for the various treatments by mixing the partitioned provisions with the selected pesticide at the desired concentration while the control is pesticide-free. After forming pellets from it, we poked a ~1mm-depth hole on each provision by the probe and seeker (Home Science Tools, MT) to fit the egg in (Fig. S3).

*Osmia* eggs were obtained by opening the paper straws to expose the bee eggs on top of the provisions. A bent spatula (Microslide Tool Set BioQuip Products Inc., CA) was used to move the egg from its own provision and place it on the prepared and treated provision. All eggs were allowed to hatch on the artificial pollen pellets in the lab at 25°C, RH 70%.

### Treatments

Pesticides selected for evaluation were chosen because they are commonly used plant systemic pesticides used during pre-bloom to control pests in apple orchards in northeastern United States (Table S1).

Acetamiprid (Assail 30SG, United Phosphorus Inc., King of Prussia, PA) is a member of the neonicotinoid class of insecticides that were introduced in the 1990s to replace the organophosphate and carbamate insecticides that were being phased out due to the implementation of the Food Quality Protection Act. It is specifically of the cyano-substituted group that is much safer for bees than the nitro-substituted neonicotinoids^43^. The current apple pesticide product label allows it to be used anytime bees are not actively foraging, including apple bloom. Although much less systemic than other members of the neonicotinoids, it has still been found in the nectar and pollen of apple blossoms sprayed five days before bloom^16^.

Flonicamid (Beleaf 50SG, FMC Corporation, Philadelphia, PA) is a selective plant systemic insecticide that intoxicates mostly through ingestion of plant sap, and although it has been found in the nectar and pollen from pre-bloom sprays^16^, it is rated by the US-EPA as practically non-toxic to bees (Minnesota Department of Agriculture). In apple, it is used primarily for the control of insecticide-resistant aphids pre-bloom^45^.

Sulfoxaflor (Closer 2SC, Dow AgroSciences LLC, Indianapolis, IN) was introduced in 2015 as a product safer to bees than the functionally similar neonicotinoids it replaced. It was immediately banned by the US Environmental Protection Agency because of those similarities and concerns about bee toxicity but has been recently re-introduced in apple and allowed to be used three days before bloom and at petal fall where it could impact pollinators.

Dodine (Syllit 3.4FL, Arysta LifeScience Benelux, Ougree, Belgium) is a foliar-applied fungicide heavily used in apple production worldwide since its commercial introduction in 1957. Because it is broad-spectrum with multiple modes of action, apple scab (*Venturia inaequalis*), resistant to other fungicides, has no known resistance to Dodine. It is used as a pre-bloom rotation partner to protect trees against this highly resistant fungal pathogen. Dodine is also plant systemic and has been found in the nectar and pollen from pre-bloom sprays^16^.

We used field-realistic concentrations of these compounds found in the pollen from apple trees treated at the same rates, methods of application, and timings as local growers^16^. Pesticide concentrations were thoroughly mixed with homogenized provisions during pellet preparation. In addition to the field-realistic (1X) rate, we also tested a 1/10X rate and a 10X to bracket possible field application scenarios used by growers in orchards that might overdose or underdose current spray recommendations found on the pesticide product label or by crop consultants (after Abbott et al. 2008)^46^. A single egg was placed on the provisions in each well container. Six replications of eight individuals were observed per pesticide treatment and control for a total of 624 individuals.

### Observation and data collection

Daily observations were made of all individuals from egg hatch until cocoon completion. The stages most easily observed were eggs, 1^st^ instar (feeding inside egg chorion), 2^nd^ instar (starting to feed on provision), 5^th^ instar (beginning defecation), initiation of cocoon spinning, and cocoon completion (Fig. S3). Growth rate and development time were assessed. Larval weight was collected daily only on the 5^th^ instars until the initiation of cocoon spinning. Pre-tests had shown that earlier instars were too delicate for this type of handling and caused excessive mortality. The net weight gain during the 5^th^ instar was calculated by deducting the weight at the initiation of cocoon spinning (pre-pupae) by the weight on the first day of the 5^th^ instar. Here we compared the average weight gain rate for the first 10 days of the 5^th^ instar because most larvae in the control started to spin cocoons on their 11^th^ day.

After the cocoon spinning was completed, feces (meconium) were cleaned from the cocoon surface, and they were transferred into Parr 3601 Gelatin Capsules (Parr Instrument Co., IL). Each capsule contained one cocoon and was numbered to keep track until emergence (Fig. S3). All cocoons were kept at 25°C, RH 70% until August 15^th^ 2017, when the temperature was gradually decreased by 5°C at the 1^st^ and 15^th^ of each month until the end of October to simulate Pennsylvania field temperatures. Cocoons were kept at −2°C from November 2017 to the end of January 2018 to provide wintering conditions for the dormant adults inside the cocoons. In February 2018, we adjusted the temperature to 2°C. Relative humidity was maintained at 70% during the whole process. At the end of March 2018 when the chilling requirements had been met^34^, we extracted adults from cocoons and assessed their weights. These temperature adjustments were utilized based on our own experiences and previous studies on the biology of *O. cornifrons*^34,47–50^.

### Statistical analysis

The overall treatment effect for all pesticide treatments was analyzed using one-way ANOVA (Minitab v.19, State College, PA, USA). The one-way ANOVA test was also used to investigate differences in development of the *O. cornifrons* larvae. Developmental differences among treatments were assessed in terms of pre-pupal development time and larval weight of 5^th^ instar. We used Fisher’s LSD comparison test for post-hoc analysis when there was a significant effect of pesticide treatment on development speed, including days required to cocoon spinning initiation, the net weight gain of 5^th^ instar, and the body weight of adult bee after emergence. Post-hoc analyses using Fisher’s LSD test contrast were also conducted when there was significant effect of pesticide treatment on the survivorship to *O. cornifrons* adult and larval abnormal defecation. In addition, same statistical procedure was used to analyze the overall survival of bees since starting the experiment until adult emergence and then assessing the adult body weight female and male bees separately. Significance level for all statistical tests was set at *α* = 0.05. Mean and standard deviations for the weight gain rate for the first 10 days of 5^th^ instar were calculated using Minitab statistical package.

## Data availability and Code availability

The datasets generated during and/or analyzed during the current study are available from the corresponding authors on reasonable request.

## Acknowledgements

This research work was partially supported by USDA-SCRI grant (PEN04398, D.J.B. and E.G.R., PDs) USDA-NIFA SCRI CAPS grant (#2012-51181-20105, R. Isaacs, PD). During this period, D.J.B. was supported by USDA-NIFA (Hatch #1011647), and N.J. was supported by the USDA NIFA (Hatch #1009954, 1024510, 1018070). Any opinions, findings, conclusions, or recommendations expressed in this publication are those of the authors and do not necessarily reflect the view of the U.S. Department of Agriculture or affiliated institutions.

## Author contributions

D.B., E.R., N.P. and N.J. conceptualized the study. N. P. developed the laboratory rearing method, and conducted bioassays with inputs from D.B., E.R., N.J., M.L., K.P. N.P. analyzed the data and developed visual graphics under the guidance of D.B., E.R. and N.J. N.P. prepared the first draft of manuscript with contributions from D.B., E.R., N.J., M.L., K.P. All authors edited/reviewed and approved the manuscript.

## Additional Information

### Supplementary Information

The online version contains supplementary material.

**Correspondence and requests for materials** should be addressed to N.P., N. K. J. or D.J.B.

## Supplementary Figures and Tables

**Figure S1.**
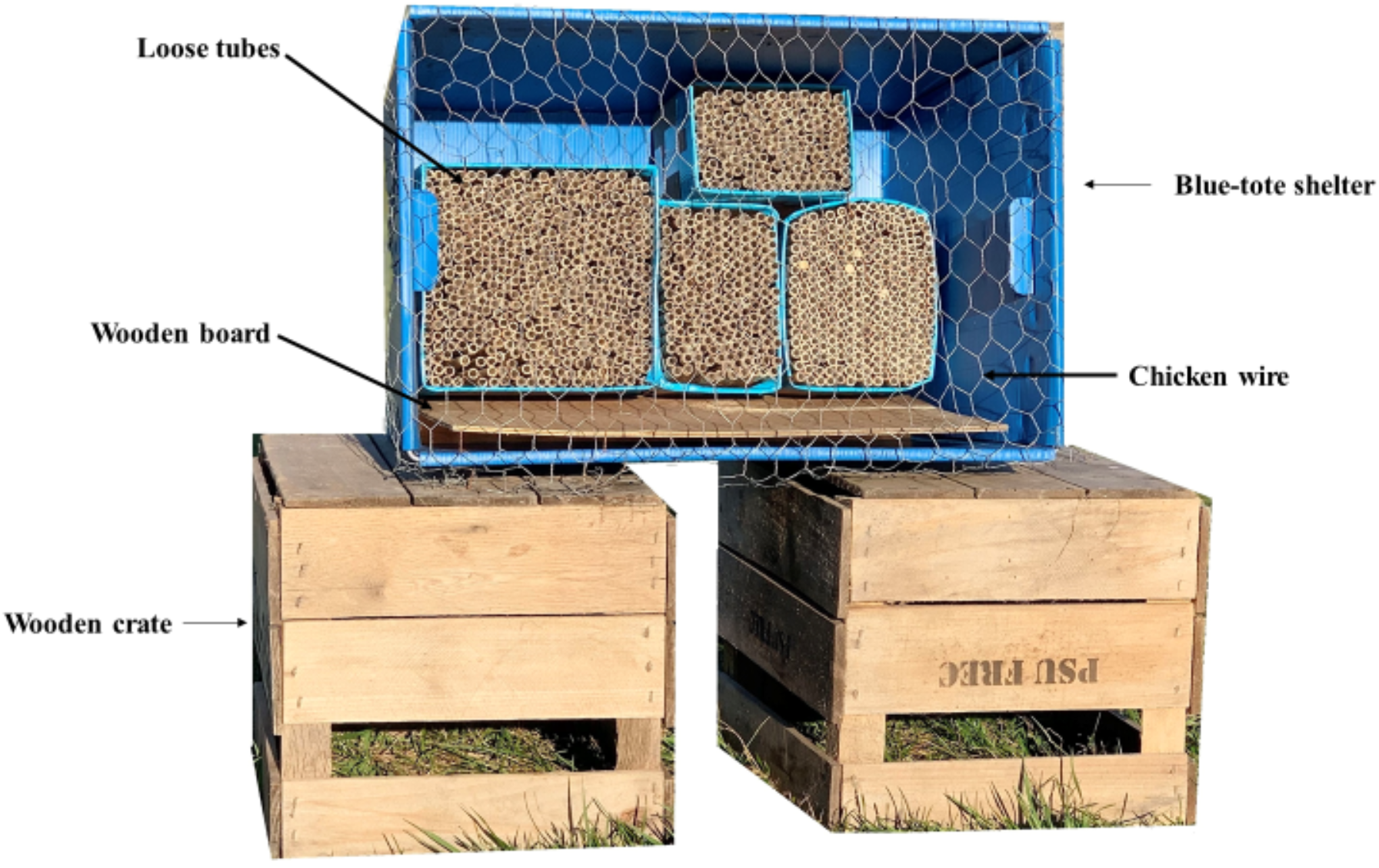
Japanese orchard bee trap nests used in the study.

**Figure S2.**
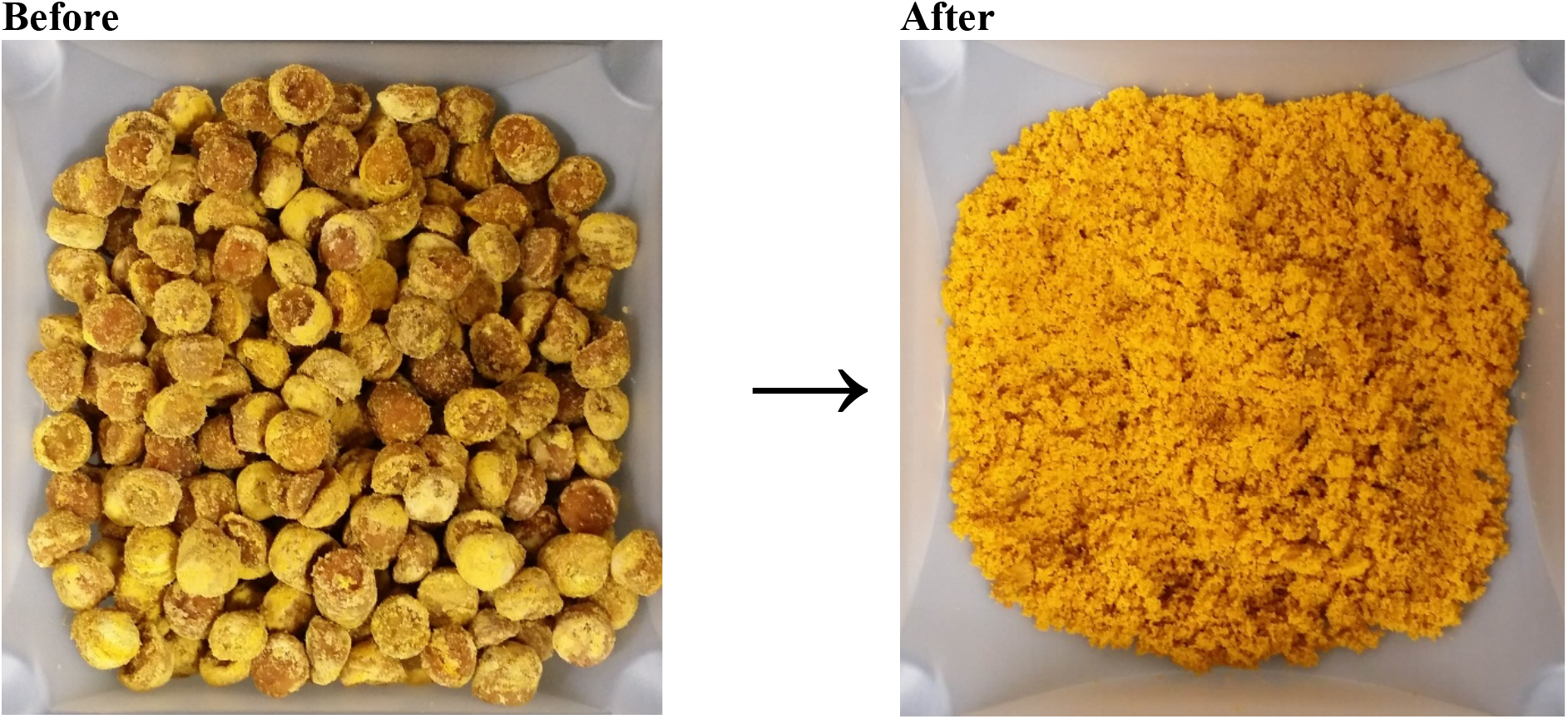
Pollen provisions before and after being homogenized by a mixer.

**Figure S3.**
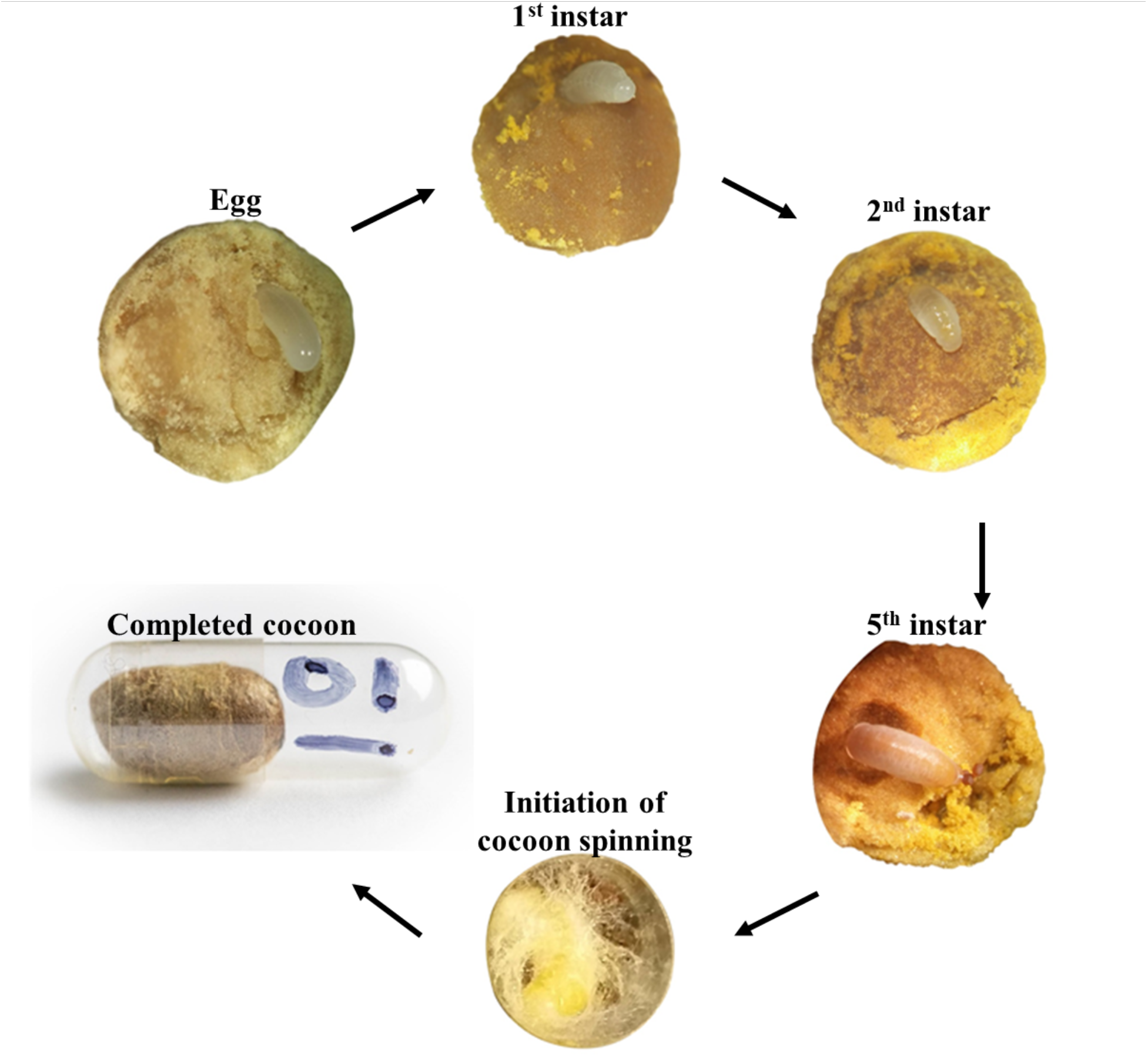
Development stages of the Japanese orchard bee (*Osmia cornifrons*) with homogenized provisions in experimental cells.

**Table S1:**
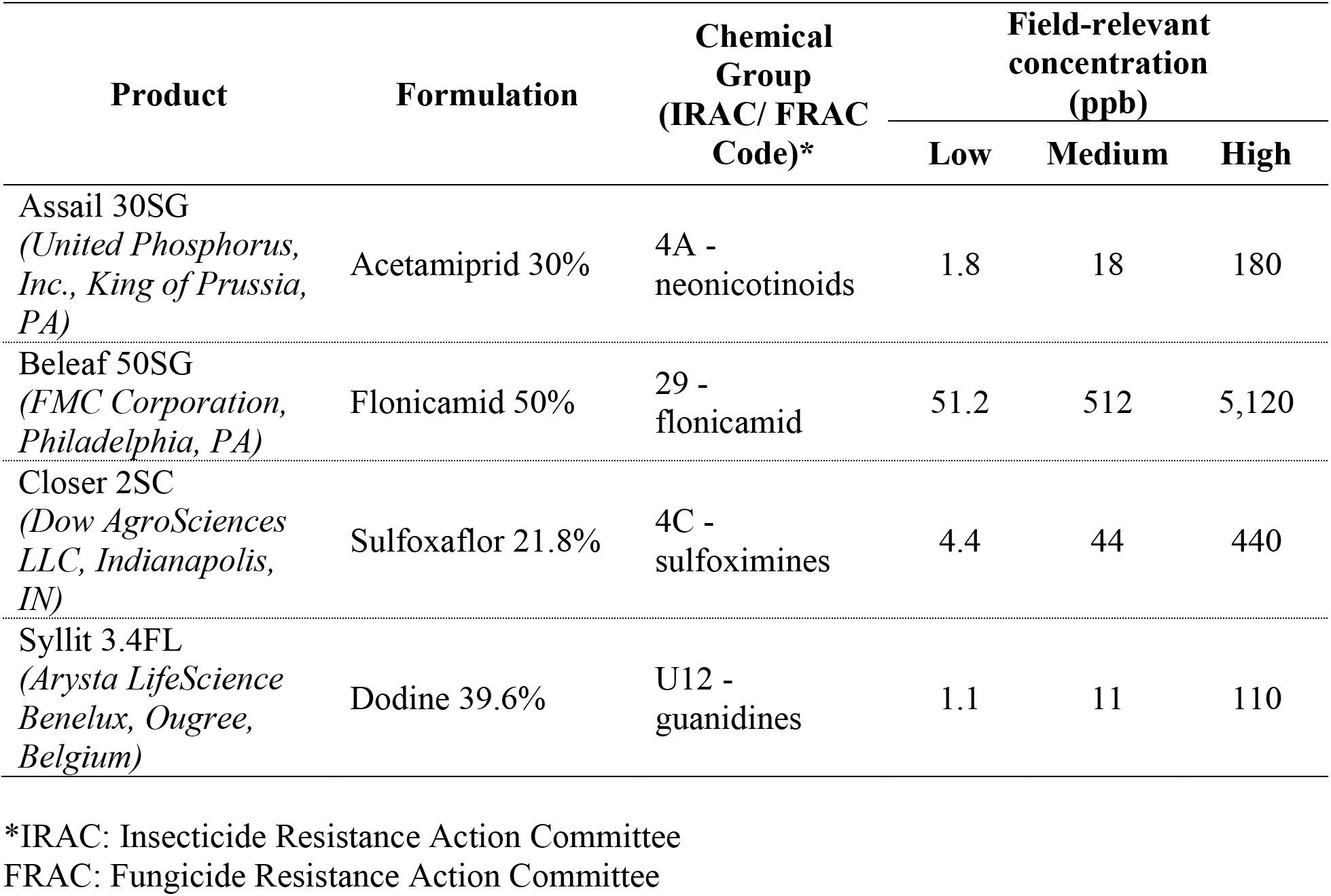
Pesticides used in larval bioassays. The commercial name, common name, and mode of action group for each pesticide are given; Chosen field-relevant concentrations (in ppb) are Low (1/10X), Medium (1X), and High (10X) concentration exposure.

