## Supplementary figures and tables for "Systemic pesticides in a solitary bee pollen food store affect larval development and increase pupal mortality"

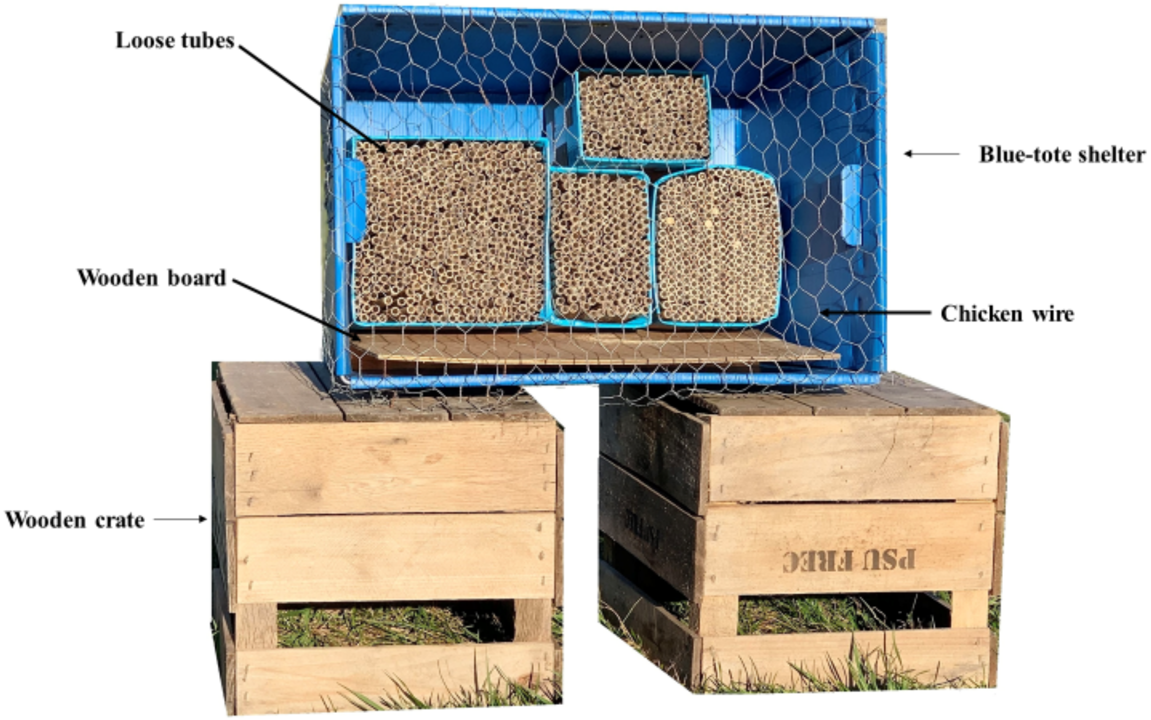


**Figure S1. Japanese orchard bee trap nests used in the study.**

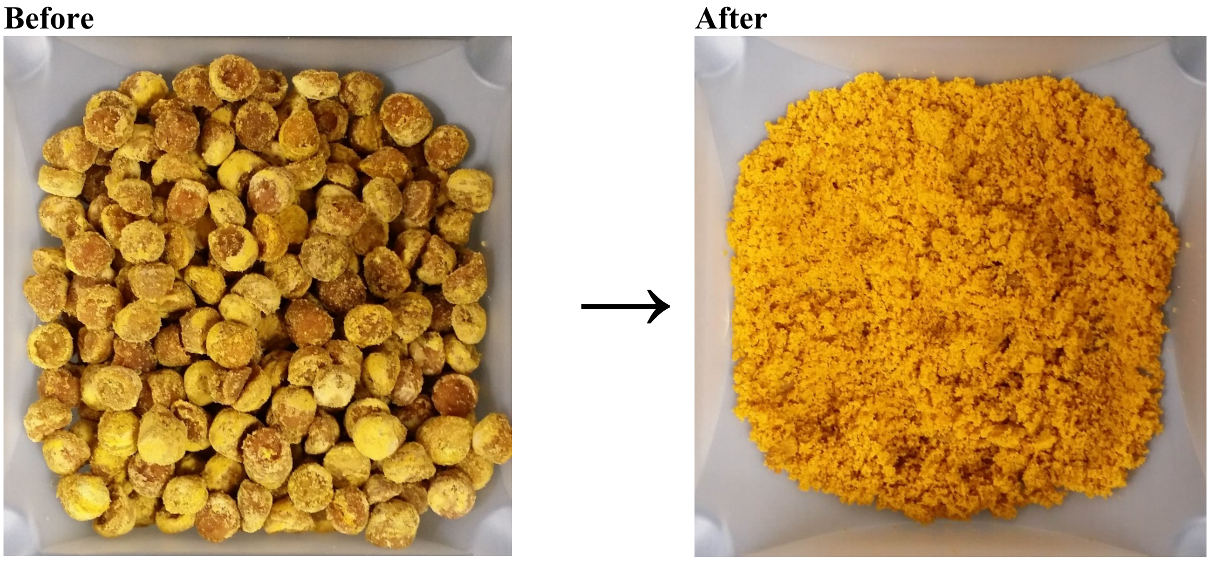


**Figure S2. Pollen provisions before and after being homogenized by a mixer.**

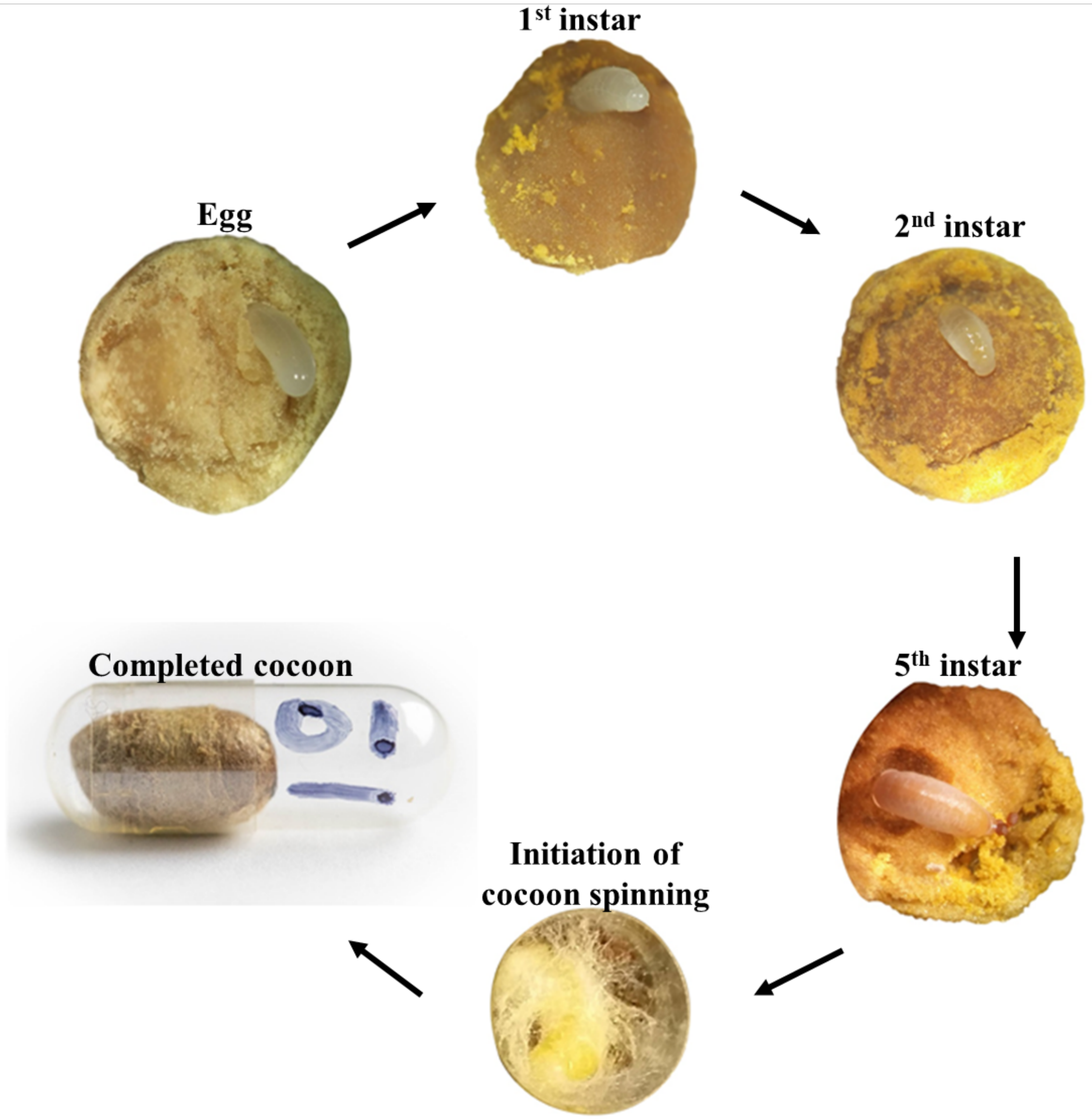


**Figure S3. Development stages of the Japanese orchard bee (*Osmia cornifrons*) with homogenized provisions in experimental cells.**

**Table S1: Pesticides used in larval bioassays.**
The commercial name, common name, and mode of action group for each pesticide are given; Chosen field-relevant concentrations (in ppb) are Low (1/10X), Medium (1X), and High (10X) concentration exposure.

| Product | Formulation | Chemical Group (IRAC/ FRAC Code)* | Field-relevant concentration (ppb) | | |
|---|---|---|---|---|---|
| | | | Low | Medium | High |
| Assail 30SG *(United Phosphorus, Inc., King of Prussia, PA)* | Acetamiprid 30% | 4A - neonicotinoids | 1.8 | 18 | 180 |
| Beleaf 50SG *(FMC Corporation, Philadelphia, PA)* | Flonicamid 50% | 29 - flonicamid | 51.2 | 512 | 5,120 |
| Closer 2SC *(Dow AgroSciences LLC, Indianapolis, IN)* | Sulfoxaflor 21.8% | 4C - sulfoximines | 4.4 | 44 | 440 |
| Syllit 3.4FL *(Arysta LifeScience Benelux, Ougree, Belgium)* | Dodine 39.6% | U12 - guanidines | 1.1 | 11 | 110 |

*IRAC: Insecticide Resistance Action Committee
FRAC: Fungicide Resistance Action Committee
